# Host species determine symbiotic community composition in Antarctic sponges (Porifera: Demospongiae)

**DOI:** 10.1101/2020.03.09.983221

**Authors:** Oriol Sacristán-Soriano, Natalia Pérez Criado, Conxita Avila

## Abstract

The microbiota of four Antarctic sponges, *Dendrilla antarctica, Sphaerotylus antarcticus, Mycale acerata*, and *Hemigellius pilosus*, collected at two South Shetland Islands and at two locations in the Antarctic Peninsula separated by ca. 670 km, were analyzed together with surrounding seawater. We used high throughput sequencing of the V4 region of the 16S rRNA gene common to Bacteria and Archaea to investigate the microbial diversity and community composition. Our study reveals that sponge-associated prokaryote communities are consistently detected within a particular sponge species regardless of the collection site. Their community structure and composition are typical of low microbial abundance (LMA) sponges. We conclude that prokaryote communities from Antarctic sponges are less diverse and differ in their composition compared to those in the water column. Microbiome analysis indicates that Antarctic sponges harbor a strict core consisting of seven OTUs, and a small variable community comprising several tens of OTUs. Two abundant prokaryotes from the variable microbiota that are affiliated to the archaeal and bacterial phyla Thaumarchaeota and Nitrospirae may be involved in the sponge nitrification process and might be relevant components of the nitrogen cycling in Antarctica. The likely generalist nature of dominant microbes and the host-specific structure of symbiont communities suggest that these Antarctic sponges represent different ecological niches for particular microbial enrichments.

## Introduction

Sponges (phylum Porifera) are sessile organisms widely distributed from the tropics to the poles (Hooper and Van Soest 2004), and ecologically important constituents of benthic environments from shallow to deep waters (Bell 2008). Sponges form symbioses with diverse and metabolically active microorganisms that make valuable contributions to many aspects of the sponge’s physiology and ecology (Taylor et al. 2007). For that reason, the concept ‘sponge holobiont’ was introduced to refer to the sponge host and the consortium of bacteria, archaea, algae, fungi, and viruses that reside within it (Webster and Taylor 2012). Marine sponge-associated prokaryotic communities have been widely studied, being reported to be highly complex (Thomas et al. 2016). Sponges host (even at low relative abundances) over 60 bacterial and 4 archaeal phyla (Reveillaud et al. 2014; Thomas et al. 2016; Moitinho-Silva et al. 2017a). Despite the continuous flux of seawater through their canal system, sponges are able to maintain a specific microbial composition remarkably different from the ambient seawater (Thomas et al. 2016; Hill and Sacristán-Soriano 2017). Furthermore, these associations appear to be host-specific and stable under different environmental conditions (Hentschel et al. 2002; Lee et al. 2011; Erwin et al. 2012b; Schmitt et al. 2012; Pita et al. 2013; Reveillaud et al. 2014). Although sponge-microbial interactions seem to be consistent over geographic regions, there are some apparent geographical gaps in the study of host-associated prokaryotic assemblages.

Antarctic marine habitats are characterized by their uniqueness and almost intact virginity that experience extreme environmental conditions and a marked seasonality. The geographical isolation and the cyclical sea-ice formation make these ecosystems largely unexplored. Sponges are also important components of marine benthic communities in Antarctica and play key roles in community structure and nutrition cycling, also providing microhabitats for other invertebrates (McClintock et al. 2005; Angulo-Preckler et al. 2018). Few studies have examined the microbial diversity present in Antarctic marine sponges (Lo Giudice et al. 2019). There have been several approximations to unravel the composition of Antarctic microbial communities associated to sponges, but primarily focused on eukaryotic microorganisms (Bavestrello et al. 2000; Cerrano et al. 2000; Cerrano et al. 2004; Henríquez et al. 2014). However, the most comprehensive studies describing microbial communities associated to Antarctic sponges using classic and high throughput sequencing described a total prokaryotic composition of 26 bacterial and 3 archaeal phyla and the presence of 8 eukaryotic groups (Webster et al. 2004; Rodríguez-Marconi et al. 2015; Cárdenas et al. 2018; Steinert et al. 2019; Cárdenas et al. 2019). Recently, functional metagenomics has been used to characterize the community composition and metabolic potential of microbiomes of two Antarctic sponges (Moreno-Pino et al. 2020). In the Bacteria domain those assemblages primarily clustered within the Gamma and Alphaproteobacteria followed by the Bacteroidetes phylum. In the Archaea domain, Crenarchaeota and Thaumarchaeota representatives were mostly associated to Antarctic sponges. Within the Eukarya domain, fungi were predominately in association with sponges in Antarctica followed by diatoms and dinoflagellates (Lo Giudice et al. 2019).

Since the rapid development in massive sequencing methodologies, allowing for a more complete characterization of the sponge microbiomes, the specificity of the associated microorganisms is under debate. Several bacterial taxa have been reported as sponge-specific bacteria (i.e., bacterial lineages found only in sponges and not in ambient seawater or sediments) (Taylor et al. 2007). However, other studies have showed that several bacterial taxa thought to be specific to sponges also occur in other habitats, such as seawater, sediment, and other hosts (Simister et al. 2012). Therefore, it would be preferable to refer to the associated microbial partners with the term ‘sponge-enriched’ or ‘host-enriched’ (Moitinho-Silva et al. 2014).

In the present study, we assess and compare prokaryote communities from four sponge species and the surrounding seawater collected in two South Shetland Islands and at two locations in the Antarctic Peninsula separated by ca. 670 km. We sought to answer the following questions: 1) How is the diversity and microbial community composition associated to Antarctic marine sponges, compared to the surrounding seawater? 2) Is there a core-microbiome associated to them? 3) Are these communities host-specific and consistent over a geographic scale? This is one of the first reports that used high throughput sequencing to unravel the composition and diversity of symbiotic microbial communities associated to Antarctic marine sponges.

## Materials and methods

### Sample collection

During the austral summer 2016, four sponge species were collected at different locations from two South Shetland Islands and the Antarctic Peninsula (Table 1). Replicate seawater samples (n = 3, 2 l samples) were aseptically collected adjacent to the sampled sponges from all locations. Sponges were processed after collection. A sample from each specimen was taken with a sterile scalpel and rinsed several times in 0.22 µm-filtered seawater to discard loose attached microorganisms. Seawater samples were passed through polycarbonate 5 µm and 0.22 µm filters (sequentially; MilliporeSigma, Burlington, MA, USA), and the contents on the 0.22 µm filters were used to examine the ambient bacterioplankton communities. All samples were preserved in RNAlater until further use.

**Table 1.** Samples of healthy specimens of *Dendrilla antarctica* (Topsent, 1905), *Sphaerotylus antarcticus* (Kirkpatrick, 1907), *Mycale (Oxymycale) acerata* (Kirkpatrick, 1907), and *Hemigellius pilosus* (Kirkpatrick, 1907) collected at 15 to 20 m depth from two South Shetland Islands (Deception and Half Moon Islands) and the Antarctic Peninsula (Rothera and O’Higgins Research Stations).

| Host species | Individuals<br>(N) | Location | Coordinates |
| --- | --- | --- | --- |
| *Dendrilla antarctica* | 4 | Whalers Bay, Deception Island | -62.984002, -60.562240 |
| *Dendrilla antarctica* | 4 | Rothera Research Station | -67.565397, -68.118247 |
| *Dendrilla antarctica* | 4 | Bernardo O'Higgins Research Station | -63.320612, -57.905138 |
| *Dendrilla antarctica* | 4 | Half Moon Island | -62.593079, -59.906964 |
| *Sphaerotylus antarcticus* | 4 | Whalers Bay, Deception Island | -62.984002, -60.562240 |
| *Sphaerotylus antarcticus* | 4 | Rothera Research Station | -67.565397, -68.118247 |
| *Sphaerotylus antarcticus* | 2 | Bernardo O'Higgins Research Station | -63.320612, -57.905138 |
| *Sphaerotylus antarcticus* | 3 | Half Moon Island | -62.593079, -59.906964 |
| *Mycale (Oxymycale) acerata* | 4 | Whalers Bay, Deception Island | -62.984002, -60.562240 |
| *Hemigellius pilosus* | 4 | Whalers Bay, Deception Island | -62.984002, -60.562240 |

### 16S rRNA gene (V4) sequence clone libraries

We used the sponges and seawater collected from Deception Island to generate these libraries. DNA was extracted using the DNeasy PowerSoil kit (Qiagen, Germantown, MD, USA) following standard protocols of the Earth Microbiome Project (http://press.igsb.anl.gov/earthmicrobiome/emp-standard-protocols/dna-extraction-protocol/). DNA was PCR amplified with the AmpliTaq Gold 360 Master Mix (Applied Biosystems) and the universal bacterial/archaeal forward and reverse primers 515fb and 806rb (Caporaso et al. 2011; Apprill et al. 2015). Three separate reactions were conducted per each sample. The thermocycler profile consisted of an initial denaturation step at 95 °C for 10 min; 31 cycles of 95°C for 45s, 51°C for 60s, and 72°C for 90s with a final elongation step at 72°C for 10 min. Equimolar concentrations of all individuals of the same species were pooled and purified using the GeneClean^®^ Turbo Kit (MP Biomedicals). Purified PCR products (ca. 10 ng) were ligated into plasmids using the pGEM^®^-T Easy Vector System (Promega).

Individual clones were PCR-screened using vector primers, and clones with approximately 250-bp inserts were purified and sequenced at Scientific and Technological Centers, Universitat de Barcelona (CCiT-UB). Bidirectional sequencing reactions were performed for all clones using vector primers. Raw sequence data were processed in Geneious (v8.1.8; Drummond et al. 2010), and low-quality sequencing reads were discarded. Representative clone sequences were deposited in NCBI GenBank database under the accession numbers MN032619-MN032636. To determine whether the cultured isolates were also recovered by the high-throughput sequencing techniques, we performed a local blast of the isolates against our 16S microbiome sequencing data (NCBI-BLAST-2.7.1+).

### Microbiome analysis

DNA extracts were submitted to Molecular Research LP (www.mrdnalab.com, Shallowater, TX, USA) for amplification, library construction and multiplexed sequencing of partial (V4) 16S rRNA gene sequences on an Illumina MiSeq platform. The HotStarTaq Plus Master Mix kit (Qiagen) was used for PCR amplifications using DNA extracts as templates with universal bacterial/archaeal forward and reverse primers 515fb and 806rb (See Cloning). To barcode samples, a multiplex identifier barcode was attached to the forward primer. The thermocycler profile consisted of an initial denaturation step at 94 °C for 3 min; 28 cycles of 94°C for 30s, 53°C for 40s, and 72°C for 1 min with a final elongation step at 72°C for 5 min. Equimolar concentrations of samples were pooled and purified using Agencourt Ampure XP beads (Beckman Coulter) to prepare DNA library by following Illumina TruSeq DNA library preparation protocol. Sequencing was then performed according to manufacturer’s guidelines on an Illumina MiSeq. Illumina sequence data were deposited in NCBI SRA under the accession number PRJNA548273.

As described previously (Thomas et al. 2016), Illumina sequence reads were processed in mothur v1.39.5 (Schloss et al. 2009). Briefly, raw reads were demultiplexed, forward and reverse reads were then joined, and sequences <200bp and/or with ambiguous base calls were removed. Sequences were aligned to the SILVA database (release 128, non-redundant, mothur-formatted), trimmed to the V4 region, and screened for chimeras and errors. A naïve Bayesian classifier and Greengenes taxonomy (August 2013 release, mothur-formatted) was used to aid in the removal of non-target sequences (e.g., chloroplasts, mitochondria). We used the SILVA database (release 132, non-redundant, mothur-formatted) for final taxonomic assignment. The resulting high-quality sequences were clustered into operational taxonomic units (OTUs) defined by clustering at 3% divergence and singletons were removed. We used rarefaction curves (mothur v1.39.5) to plot the OTUs observed as a function of sequencing depth. To avoid artifacts of varied sampling depth on subsequent diversity calculations, each sequence dataset was subsampled to the lowest read count (mothur v1.39.5). To place the determined OTUs into a greater context, these were compared to the database of the sponge EMP project (Moitinho-Silva et al. 2017a) using local BLAST searches (NCBI-BLAST-2.7.1+).

### Community-level analysis

To compare bacterial community profiles, nonmetric multi-dimensional scaling (nMDS) plots of Bray-Curtis similarity matrices were constructed with mothur (v1.39.5) and R (version 3.4.3; ggplot2 package) from square-root transformed OTU relative abundance data. We also constructed bubble charts in R (version 3.4.3; ggplot2 package) from OTU relative abundances to plot community dissimilarities among locations. Significant differences among sponge species and ambient seawater were assessed using a one-way permutational multivariate analysis of variance (PERMANOVA), with the factor source (all sponge species vs. seawater). Significant differences among sponge species were assessed with a one-way PERMANOVA, with the factor source (*D. antarctica, S. antarcticus, M. acerata*, and *H. pilosus*). Differences between sponge species and locations were assessed using a two-way PERMANOVA, with the factors source (*D. antarctica* and *S. antarcticus*), location (Deception Island, Rothera, Half Moon Island, O’Higgins) and an interaction term. Pairwise comparisons were subsequently conducted for all significant PERMANOVA results. Permutational multivariate analysis of dispersion (PERMDISP) was used to detect differences in homogeneity (dispersion) among groups for all significant PERMANOVA outcomes. All multivariate statistics were performed in R (version 3.4.3; with adonis2 and betadisper functions from vegan v2.5-6 package).

We calculated three indices of alpha diversity in mothur v1.39.5 (Schloss et al. 2009) to evaluate community richness and evenness: observed OTU richness, the Simpson index of evenness and the inverse of Simpson index of diversity. One-way analyses of variance (ANOVA) was used to detect differences in diversity metrics among the species from Deception Island (*D. antarctica, S. antarcticus, M. acerata*, and *H. pilosus*). Two-way analysis of variance (ANOVA) was used to detect differences in diversity metrics by the factors source (*D. antarctica* and *S. antarcticus*), location (sampling sites) and an interaction term, followed by pairwise comparisons for any significant factor. All data that did not meet the statistical assumptions was transformed accordingly (log-transformation for inverse of Simpson index). The univariate statistics were performed in R (version 3.4.3; Anova function from car package).

### OTU-level analysis

We were interested in OTUs that were abundant (i.e., >0.1% relative abundance) and widespread among sponge host individuals (i.e., 90% prevalence), so we performed a core microbiome analysis in R (version 3.5.3, package Microbiome). We also analyzed the dataset for patterns in relative abundances of OTUs within categories (e.g., sponge vs. seawater, among locations). For this purpose, we removed from the dataset rare OTUs (<0.1% relative abundance) and OTUs with a low incidence across samples (detected in <2 samples). We used the Mann-Whitney-U test (or Wilcoxon rank sum test) with FDR p-value correction to identify significantly different patterns in OTU relative abundance among hosts and life stages using QIIME (Caporaso et al. 2010).

## Results

### Microbiome composition associated to Antarctic sponges

The V4 region of the 16S rRNA gene was sequenced on an Illumina MiSeq platform and a total of 4,398,237 reads were obtained after denoising and quality filtering with a library depth ranging from 39,883 to 154,480 reads. As we had 4 replicates per species and location in most of the cases, we discarded those samples (n = 5) with the lowest number of reads (≤51,407) to have at least 3 replicates per sponge and site. To avoid artifacts of varied sampling depth, we rarefied our libraries to the lowest read count after removing the previous samples from the dataset (n = 52,637; Suppl. Fig. S1). Twenty-eight bacterial and 3 archaeal phyla were detected in the 11,187 OTUs recovered from seawater and sponge samples, which were predominantly affiliated to the phyla Proteobacteria and Bacteroidetes (Suppl. Fig S2). Of these, 4,619 OTUs were recovered from *D. antarctica*, 3,438 OTUs from *S. antarcticus*, 1,381 OTUs from *M. acerata*, and 1,490 OTUs from *H. pilosus* Seawater exhibited greater richness with 6,071 OTUs, 2,511 of which were shared with either *D. antarctica* or *S. antarcticus* or with both species, and 1,240 were shared with *M. acerata* and/or *H. pilosus* (Suppl. Fig. S3). Seawater from Rothera, and especially from Deception Island contained the lowest amounts of Gammaproteobacteria.

The taxonomic composition of microbial communities recovered from ambient seawater, and from *S. antarcticus, M. acerata* and *H. pilosus* sponge hosts, were significantly different, while *D. antarctica* presented a microbial content quite similar to what we found in seawater (Fig. 1). Firstly, the microbial communities harbored by the first three species were enriched for Gammaproteobacteria (>60%, >80%, >65% of the microbial community on average, respectively) and Thaumarchaeota (>20% in *S. antarcticus* and *H. pilosus*, >8% in *M. acerata*) compared to seawater (<42% and <0.5%, respectively). Comparatively, *D. antarctica* hosted less Gammaproteobacteria (<54%) and its associated Thaumarchaeota were almost absent (<0.5%). Secondly, microbial communities in the first three sponges were depleted in members of Alphaproteobacteria (<5% in all three species) and Bacteroidetes (<8% in all three species) compared to seawater (>24% and >30%, respectively) and *D. antarctica* (>26% and >17%, respectively). The inter-individual variability of the taxonomic composition depends on the host. While *H. pilosus* and *S. antarcticus* harbored a quite stable microbial signature, *D. antarctica* exhibited greater inter-individual variability and one of the specimens of *M. acerata* showed an enrichment for Thaumarchaeota (Fig. 1).

**Figure 1.**
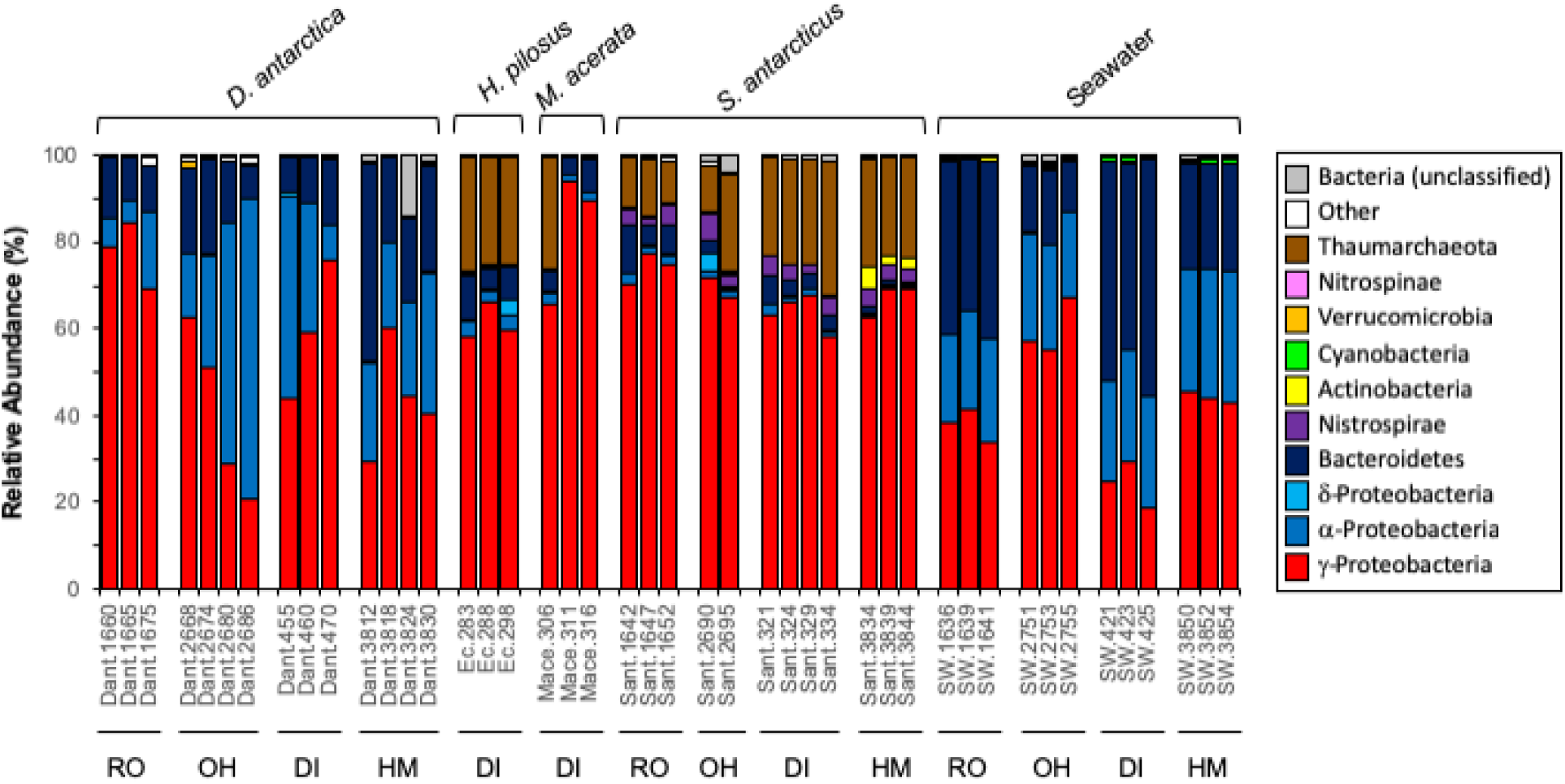
Taxonomic composition of bacterial communities in *Dendrilla antarctica, Sphaerotylus antarcticus, Mycale acerata, Hemigellius pilosus* and surrounding seawater from O’Higgins (OH), Half Moon Island (HM), Deception Island (DI) and Rothera (RO).

### Differences within and between sponge-associated and seawater microbial communities

Statistically significant differences in community structure (PERMANOVA) were detected among *S. antarcticus, M. acerata, H. pilosus* and *D. antarctica* and seawater microbes (F_4,39_ = 10.563; P = 0.001). Symbiont communities from seawater sources exhibited no overlap with sponge species in the multidimensional space, and all sponge species occupied distinct regions of the nMDS plot (Fig. 2). In addition, a significant interaction between host species (*S. antarcticus* and *D. antarctica*) and location occurred (PERMANOVA, F_3,18_ = 2.422; P = 0.008), though we could not detect significant pairwise differences in community structure after p-value correction. Dispersion analysis revealed equal variability within *S. antarcticus* and *D. antarctica* microbial communities regardless of location (P > 0.05 in all comparisons), but microbiomes of the latter species from O’Higgins were more variable (P = 0.046; Fig. 2).

**Figure 2.**
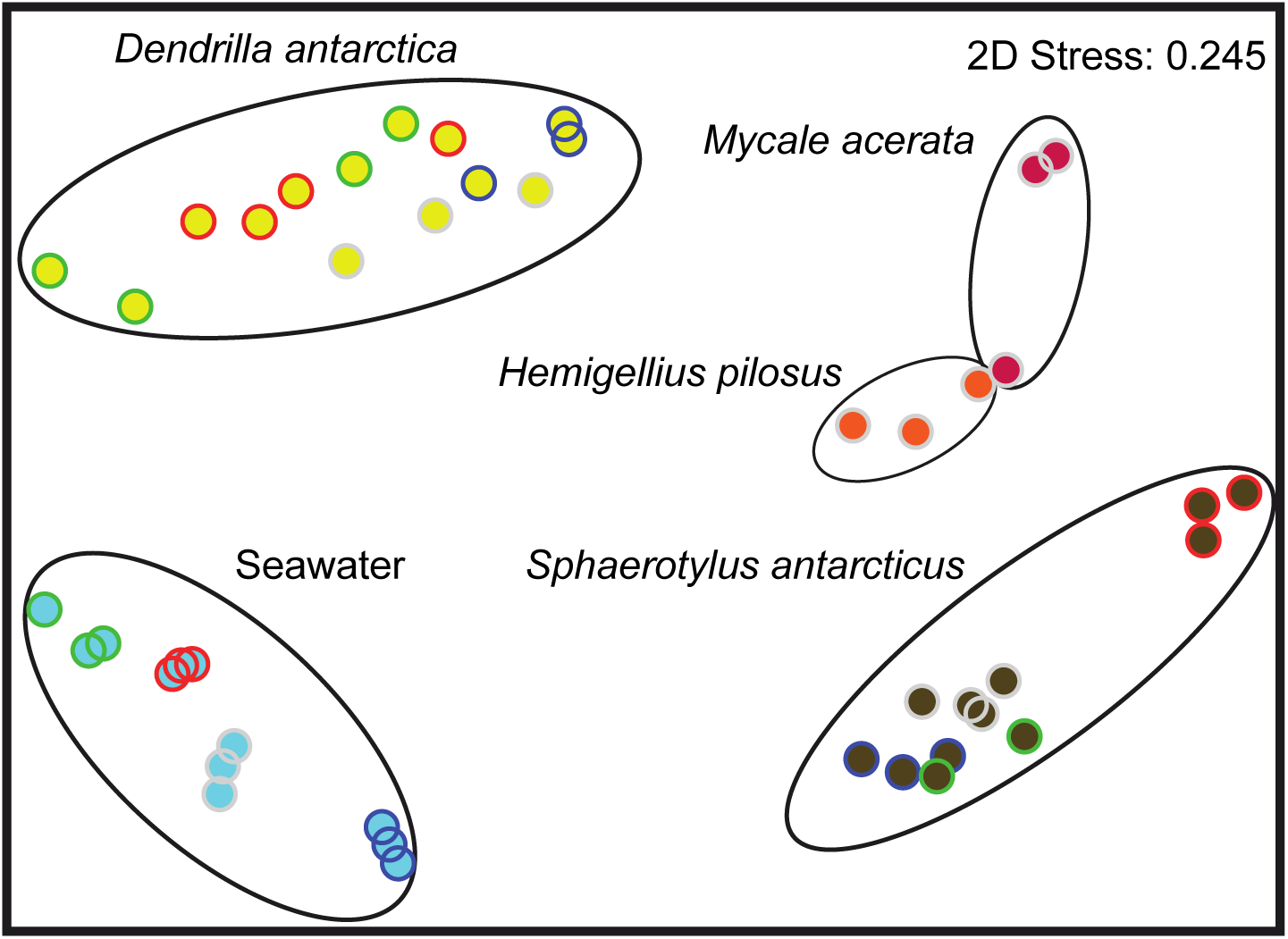
Nonmetric multi-dimensional scaling plot of microbial community structure from replicate individuals of *Dendrilla antarctica* (yellow), *Sphaerotylus antarcticus* (brown), *Mycale acerata* (red), *Hemigellius pilosus* (orange) and surrounding seawater (light blue) from O’Higgins (green circles), Half Moon Island (red circles), Deception Island (gray circles) and Rothera (blue circles). Stress value for two-dimensional ordination is shown.

Larger mean values of richness, diversity (i.e., inverse Simpson diversity index), and evenness in symbiont communities from seawater compared to host species were observed (P < 0.001 in all pairwise comparisons; Table 2). When we analyzed the sponges from Deception Island, the microbiome of all the species seemed to be equally richer and diverse with similar evenness except for the comparison between *H. pilosus* and *M. acerata*, where the former species presented more OTU richness (P = 0.048). Comparing *D. antarctica* and *S. antarcticus* from the four locations studied, a two-way ANOVA detected significant differences between hosts and locations for species richness, diversity and evenness (P < 0.03 in all cases). *D. antarctica* harbored a richer microbiome but less diverse than *S. antarcticus*. Additionally, the sponge microbial communities from Half-moon Island had greater species richness and diversity compared to those from Rothera (pairwise comparisons P < 0.05). Although there was an effect of location on the microbiome evenness, differences among pairs were not detected.

**Table 2.** Diversity estimators for microbial communities associated with seawater, *Dendrilla antarctica, Sphaerotylus antarcticus, Mycale acerata* and *Hemigellius pilosus* from O’Higgins (OH), Half Moon Island (HM), Deception Island (DI) and Rothera (RO). All values represent means (±SE).

| Source | OTU richness | Inverse Simpson's diversity | Simpson's evenness |
| --- | --- | --- | --- |
| Seawater |  |  |  |
| OH | 1718 (48.40) | 10.52 (2.31) | 0.006 (0.0007) |
| HM | 1479 (27.01) | 11.63 (0.37) | 0.008 (0.0001) |
| DI | 1143 (53.19) | 7.39 (0.44) | 0.006 (0.0002) |
| RO | 1198 (121.67) | 11.42 (0.64) | 0.010 (0.0002) |
| *D. antarctica* |  |  |  |
| OH | 900 (99.21) | 4.04 (1.23) | 0.005 (0.0014) |
| HM | 913 (78.30) | 4.85 (1.35) | 0.005 (0.0012) |
| DI | 766 (59.18) | 2.74 (0.74) | 0.004 (0.0011) |
| RO | 680 (80.91) | 2.06 (0.68) | 0.003 (0.0006) |
| *S. antarcticus* |  |  |  |
| OH | 740 (21.21) | 3.96 (0.65) | 0.005 (0.0007) |
| HM | 697 (31.97) | 5.41 (0.66) | 0.008 (0.0012) |
| DI | 669 (36.61) | 3.96 (0.39) | 0.006 (0.0005) |
| RO | 647 (67.47) | 3.74 (0.18) | 0.006 (0.0005) |
| *M. acerata* |  |  |  |
| DI | 653 (122.46) | 2.90 (2.73) | 0.004 (0.0035) |
| *H. pilosus* |  |  |  |
| DI | 784 (91.24) | 5.99 (0.55) | 0.008 (0.0008) |

The abundance of shared OTUs (n = 2,893) between sponge-associated and seawater microbial communities was calculated and just 2.2% presented relative abundances over 0.1%. Those few OTUs (n = 65) accounted for 93% and 89% of the total relative abundance of sponge-associated and seawater microbial communities, respectively, which meant that sponge-specific OTUs (64%) fell within the ‘rare biosphere’ (<0.1% relative abundance).

### Core and variable microbiome in Antarctic sponges

In addition to community-level metrics of diversity and structure, we performed a core microbiome analysis to investigate patterns in abundant and prevalent individual OTUs among sponge hosts. The strict core microbiome (i.e., with relative abundances >0.1% and present in all species) of the sponge hosts was formed by 7 OTUs accounting for 50% of total relative abundance on average (0.1% of total OTUs present in sponge hosts, 4.2% of OTUs with relative abundance >0.1% in at least one sample; Suppl. Table S1). Significant sponge enrichments in 4 core OTUs, affiliated to Gammaproteobacteria, were detected with a mean fold-change in abundance of 44.9 ± 8.0 (± SE) with respect to seawater (mean relative abundance 0.28%). Three additional sponge core OTUs, which were affiliated to the groups Alphaproteobacteria, Bacteroidetes, and Gammaproteobacteria, were more abundant in seawater communities (fold-change 14.3 ± 4.5; Suppl. Table S2). We also determined the variable community (i.e., with relative abundances >0.1% and present in at least two species) formed by 56 OTUs that represented on average 39% of the total abundance (0.7% of total OTUs present in sponges, 33.3% of OTUs with a relative abundance >0.1% in at least one sample; Suppl. Table S1). Forty-three sponge variable OTUs that had a mean fold-change in abundance of 49.5 ± 8.2 were extremely rare in seawater (mean relative abundance <0.03%). Thirteen additional sponge variable OTUs were enriched in seawater with mean relative abundance >2.7% (fold-change 13.7 ± 2.6). We comparatively determined the core microbiome of *D. antarctica* and *S. antarcticus*. Both species harbored 10 major OTUs (6 out of 7 sponge core OTUs and 4 variable OTUs) representing over 70% of the microbiome relative abundance (Suppl. Table S1). Seawater presented instead a core microbiome of 18 OTUs (including 6 sponge core OTUs) accounting for over 60% of the microbiome in relative abundance (Suppl. Table S1). Comparing the sponge species analyzed, 4 core and 17 variable OTUs seemed to be sponge-enriched for either *D. antarctica* (OTUs 1, 4, 9, 11, 13, 27, 33 and 45; cumulative 84% relative abundance; Fig. 3), *S. antarcticus* (OTUs 2, 7, 23, 31, 43, 51 and 56; 54% relative abundance; Fig. 3), *M. acerata* (OTU 6; 65% relative abundance) or *H. pilosus* (OTUs 3, 14, 19, 34 and 53; 71% relative abundance), although they were also detected in the other hosts and in seawater in lower frequencies (Suppl. Table S2). If we compared locations, neither *D. antarctica* nor *S. antarcticus* presented differences in their microbiome abundances among sampling sites (Fig. 3; Suppl. Table S2). However, OTU 2 decreased its relative abundance to 0.3% in *S. antarcticus* at Half Moon Island, whereas the proportion was over 40% on average for the rest of locations (Fig. 3).

**Figure 3.**
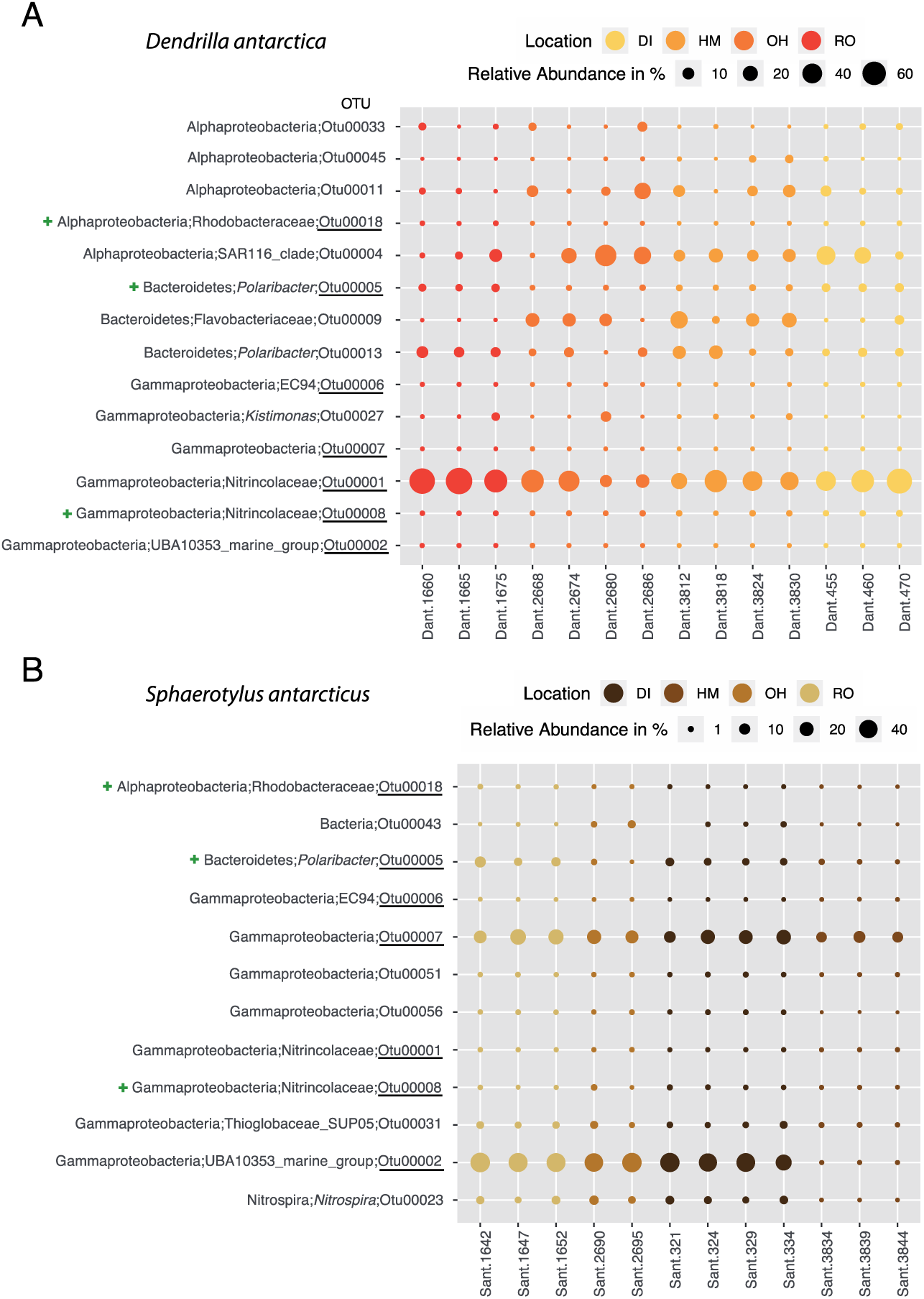
Bubble chart of core (underlined) and enriched variable OTUs of *Dendrilla antarctica* (A) and *Sphaerotylus antarcticus* (B) among locations defined at >0.1% relative abundance. OTU relative abundances are represented by the size of the bubbles (key on the top of each chart; notice the different scale between A and B). The smallest taxonomical level for each OTU is also shown. Location key: O’Higgins (OH), Half Moon Island (HM), Deception Island (DI) and Rothera (RO). We also show with a green cross those OTUs enriched in seawater samples.

From the 8,009 OTUs present in the sponges analyzed, 56.1% could be described as host-specific (i.e., present only in one sponge species), ranging from 254 OTUs in *M. acerata* to 2,230 OTUs in *D. antarctica*. However, all of these OTUs belonged to the rare biosphere associated to the sponge host with mean relative abundance below 0.1%. Although OTUs from core and variable microbial communities were detected in all the sponge species analyzed, and even in ambient seawater, the specificity for one of the hosts of some variable OTUs (38%) could be considered due to the low number of reads recovered from the other habitats (<0.02% relative abundance).

### Comparing Antarctic sponge associated microbial communities with the sponge EMP database and clone libraries from the present study

Local BLAST searches against the sponge EMP database showed that 87% of the OTUs (n = 9,782) were found among the sponge microbiome collection at high sequence similarities (Suppl. File S1). Both core and variable communities associated to Antarctic sponges had a closest relative in the sponge microbiome database with sequence identities over 97%. Only 3 variable OTUs had similarities between 93-96%. The core microbiome associated to the Antarctic sponges is also associated to other sponge hosts and habitats (Suppl. File S1).

From the clone library analysis, the V4 region of the 16S rRNA gene sequenced on a Sanger platform yielded a total of 112 sequences after denoising and quality filtering. Each library, composed by all replicates from either one sponge host or seawater, ranged from 6 to 33 sequences. All these sequences were classified into 18 clone OTUs, mostly unique in the sources analyzed (Suppl. Fig. S4). Local BLAST searches against the 16S microbiome data from this study showed that 100% of the clone OTUs were found among our microbiome collection with sequence identities over 98% (Suppl. File S2). We could recover 6 OTUs assigned to the core microbiome (>0.1% relative abundance and 90% prevalence) and 8 variable OTUs (>0.1% relative abundance and 10-90% prevalence) from sponge hosts, accounting for over 60% of the total microbiome relative abundance.

## Discussion

This work describes the bacterial and archaeal diversity and the community composition of four Antarctic sponge species, revealing that the sponges had a microbial signature different from the richer and diverse seawater community. In Antarctica, due to the difficult access to this region, only few studies have made comprehensive descriptions of microbial symbionts of marine sponges (Webster et al. 2004; Rodríguez-Marconi et al. 2015; Cárdenas et al. 2018; Steinert et al. 2019; Cárdenas et al. 2019; Moreno-Pino et al. 2020). However, this is the first study that assesses the microbial communities associated to Antarctic sponges using high throughput sequencing, considering biological replications and covering a spatial scale over 650 km.

### Diversity and taxonomic composition of microbial communities associated to Antarctic sponges

Although there is a lot of diversity to be uncovered, we have captured all abundant microbes present in Antarctic sponges and seawater (Suppl. Fig. S1). The sponge associated Bacteria/Archaea communities were less diverse than surrounding seawater as previously reported for other Antarctic sponge hosts, reinforcing the view that these sponges were composed of low microbial abundance (LMA) microbiomes (Moitinho-Silva et al. 2017b). Although Antarctic sponges displayed less diversity than the surrounding seawater, two more phyla were detected in sponges (Suppl. Fig. S5 & S6). If we discard phyla with low sequence abundances (i.e., <0.1% abundance), sponges and seawater harbored 5 bacterial and 1 archaeal phyla. Both biotopes presented Thaumarchaeota as the predominant archaeal phylum, which accumulated 10.8% of the reads in sponges while just 0.1% in seawater. With regard to bacteria, differences lay in the fact that sponges hosted bacteria of Nitrospirae and Nitrospinae phyla, while seawater had instead representatives of Cyanobacteria and Verrucomicrobia. However, all these phyla accumulated microbial abundances ranging from 0.1 to 1.4%.

Classes Gamma- and Alphaproteobacteria dominate bacterial assemblages in association with marine sponges from different biotopes (Thomas et al. 2016; Moitinho-Silva et al. 2017a; Pita et al. 2018; Cleary et al. 2019). These classes also dominate Antarctic sponges as previously reported on Antarctic marine shallow (Webster et al. 2004; Rodríguez-Marconi et al. 2015; Cárdenas et al. 2018; Cárdenas et al. 2019) and deep waters (Steinert et al. 2019) with Bacteroidetes also as important members in terms of number of OTUs recovered (n = 1,174) and in relative abundance (10%). The presence of the phylum Nitrospirae (>3% in relative abundance) in one of the sponges analyzed (*S. antarcticus*) is particularly noticeable. Members of this phylum are potentially involved in nitrification processes, specifically in nitrite oxidation (Radax et al. 2012). A previous study provided a preliminary description of the bacterial communities present in Deception Island waters, which were very different than the usual Antarctic microbial diversity (Angulo-Preckler et al. 2015). We found that Bacteroidetes dominated the seawater microbial community, probably influenced by the special conditions of this island, an active volcano with high concentrations of geothermal elements. This suggested a correlation between the environmental microbial diversity and the geochemical composition of the island. However, the sponges studied from the same area contained a different microbial signature than the environment.

The phylum Chloroflexi, and other bacterial phyla such as Actinobacteria or Acidobacteria, have been frequently found in high microbial abundance (HMA) sponges (Schmitt et al. 2011; Moitinho-Silva et al. 2017b), occasionally reaching high percentages in relative abundances (5-10%). Contrastingly, these phyla represent a much lower percentage in Antarctic sponges (0.0005-0.4%). Our results are in agreement with previous studies carried out in Antarctica (Rodríguez-Marconi et al. 2015; Cárdenas et al. 2018; Steinert et al. 2019; Moreno-Pino et al. 2020). Other widely described but less abundant phyla associated with sponges from other geographical areas are Poribacteria and PAUC34f (Moitinho-Silva et al. 2017b). These bacterial groups were not detected neither in the Antarctic sponges from the present study nor in other species previously studied in Antarctica (Rodríguez-Marconi et al. 2015; Cárdenas et al. 2018; Steinert et al. 2019; Cárdenas et al. 2019; Moreno-Pino et al. 2020). These phyla have been found overrepresented in HMA over LMA sponges and can be used as “HMA/LMA indicators” (Moitinho-Silva et al. 2017b; Glasl et al. 2018). This suggests that the microbial community structure of the sponges from the present study resemble that from LMA sponges.

Besides Bacteria, the presence of Archaea associated to marine sponges from tropical to cold waters has been previously documented (Radax et al. 2012; Jackson et al. 2013; Cardoso et al. 2013; Kennedy et al. 2014; Polónia et al. 2014; Turon and Uriz 2020). These archaeal communities were dominated by the phylum Thaumarchaeota, which can reach high relative abundances in the sponge prokaryotic community. Three of the species analyzed in this study (*S. antarcticus, M. acerata, H. pilosus*) harbored abundant populations of an OTU (∼10-25%) affiliated to *Candidatus* Nitrosopumilus. In Antarctica, the presence of Thaumarchaeota-related sequences has been reported in multiple species (Steinert et al. 2019; Moreno-Pino et al. 2020), including *M. acerata* (Webster et al. 2004; Rodríguez-Marconi et al. 2015). These taxa act as ammonia oxidizers not only in tropical reefs but also in cold environments (Radax et al. 2012; Cardoso et al. 2013; Polónia et al. 2015). Indeed, the genomic potential for ammonia oxidation was recently evidenced for *Leucetta antarctica* microbiome from Antarctica (Moreno-Pino et al. 2020). Together with the potential nitrite oxidizers detected (members of Nitrospirae), they may be considered as relevant factors in nitrogen cycling in Antarctica. However, further studies are needed to understand the functional roles these microorganisms play in Antarctic symbiosis and their contribution to the ecosystem.

### Core, variable and species-specific microbiome in Antarctic sponges compared to seawater communities

A minimal core microbial community (0.1%) was found in Antarctic sponges consisting of very few OTUs (n = 7) present in all four species but also in ambient seawater. The core prokaryotic community of sponges is rather small, pattern that has been documented from different biogeographic regions, including Antarctica (Schmitt et al. 2012; Easson and Thacker 2014; Rodríguez-Marconi et al. 2015). We also defined a small variable community (0.7%) that consisted of 56 OTUs hosted in at least two of the sponge species analyzed with relative abundances above 0.1%. Nevertheless, 43% of these variable taxa had such low abundances (<0.001% on average) in three of the species that they could be considered as species-specific OTUs. The concept of ‘sponge or host specificity’ needs to be revised as molecular techniques now allow deep sequencing of associated microbial assemblages. A taxon we thought unique of a particular biotope (e.g., sponges, a particular host), we are able now to detect it in other habitats but at much lower numbers. Thus, the term ‘sponge- or host-enriched’ has been introduced (Moitinho-Silva et al. 2014). In this regard, *D. antarctica* and *M. acerata* presented an enrichment of one of the core OTUs and *S. antarcticus* was enriched with other two core taxa representing a 45-fold change compared to seawater. The additional three sponge core OTUs were enriched in planktonic communities (Suppl. Table S2). The same enrichment pattern was found in 30% of the variable OTUs (Suppl. Table S2), which were categorized as rare (0.06% on average) in the surrounding seawater representing a 50-fold increase.

The fact that predominant OTUs were detected in all sponges from different geographic locations and also in the surrounding seawater suggests a global distribution of these microbes through the Antarctic environment. These OTUs might be adapted to their particular niches representing different ecotypes of the same microorganisms, so it is possible that they are horizontally transmitted through strong selective mechanisms (Schmitt et al. 2012; Turon et al. 2018). The majority of the sponge prokaryotic community (99%) belonged to the ‘rare biosphere’ with mean relative abundances below 0.1%, and over 50% could be described as host-specific taxa. The specificity of host-microbe associations seems to extend beyond some dominant taxa from the variable community into the rare biosphere (Reveillaud et al. 2014).

Beyond the existence of certain plasticity of microbiomes within a particular host, the dominance of a gammaproteobacterial OTU (Betaproteobacteriales) in *M. acerata* was also recently reported from another location of the Antarctic Peninsula (Cárdenas et al. 2019). This spatial stability was also documented in the present study for dominant OTUs in *D. antarctica* and *S. antarcticus* across a geographic scale <700 km. These results are in agreement with a recent study in the Caribbean that found little variations in the sponge microbiome of *Cliona delitrix* over small geographic scales (<300 km), while a considerable geographic distance impact over a large regional scale (>1,000 km) was reported (Easson et al. 2020). To date, published data support the combination of host identity, geography, and environment as the main forces determining the structure of sponge microbiomes (Webster et al. 2010; Schmitt et al. 2012; Easson et al. 2020). In Antarctica in particular, the microbial signature of sponges might be also related to a biogeographic partitioning of Southern Ocean microorganisms caused by the Polar Front, such as the deficit of Cyanobacteria in Antarctica (Wilkins et al. 2013), as evidenced in the present and previous studies (Rodríguez-Marconi et al. 2015; Cárdenas et al. 2018; Moreno-Pino et al. 2020). Since most collection efforts of sponges have so far explored tropical and temperate environments, this study contributes to expand our knowledge on sponge microbiome structure in polar waters.

### Comparison of Antarctic sponge microbiomes with clone libraries and sponge EMP database

We have demonstrated with this study that we can recover sponge core OTUs cloning the 16S rRNA gene sequence. However, if a more comprehensive and thorough analysis of the host-associated microbial assemblages is needed, high throughput sequencing techniques are required. The fact that nearly 90% of the Antarctic sponge microbiomes had a blast hit with a sequence similarity over 97% against the sponge EMP collection would represent the ubiquity of the sponge-associated microbes through species and habitats.

## Conclusion

Antarctic sponge-associated microbial communities displayed less diversity than their surrounding seawater counterparts conferring them the status of LMA sponges. Their microbial composition and structure with one or two dominant OTUs also resembled that from LMA sponges. Some abundant microbes have been related to the nitrification process and may play a central role in the nitrogen cycling in Antarctica. A global distribution of sponge-associated microbes has been documented; however, symbiont communities exhibit little uniformity in species composition or structure (Thomas et al. 2016). Antarctic sponges seem to follow the same pattern of symbiotic community organization. The core microbiomes are characterized by generalists microbes with little representation of specialists, a pattern previously described as ‘specific mix of generalists’ (Erwin et al. 2012a). Different sponge species likely represent different ecological niches for prokaryotes, each with a specific microbial community that is vertically and horizontally acquired and selectively maintained (Webster et al. 2010; Sacristán-Soriano et al. 2019). Host identity seems to be the strongest driving force in determining the composition of sponge symbiont assemblages. However, associated microbial communities could be slightly influenced by biogeography and environmental factors defining a microbial signature for a particular habitat (Kennedy et al. 2014; Rodríguez-Marconi et al. 2015; Easson et al. 2020). In future studies, the use of metagenomics (Moreno-Pino et al. 2020) and metatranscriptomics will allow the recovery of functional genes and improve our understanding of the physiological roles of Antarctic sponge-associated microbiota.

## Acknowledgements

We would like to thank BIO-Hesperides and BAE Gabriel de Castilla crews for their logistic support during the XXI Spanish Antarctic Survey. Thanks are also due to C. Leiva, P. Álvarez-Campos, G. Giribet, J. Junoy, B. Figuerola, J. Moles, C. Angulo-Preckler, and L. Cardona, for their help with sample collection. Special thanks to F.J. Cristobo for his valuable help in sample collection and taxonomic identifications. This work was funded by grants from the Spanish Government: DISTANTCOM (CTM2013-42667/ANT), BLUEBIO (CTM2016-78901/ANT), and PopCOmics project (CTM2017-88080; MCIU,AEI/FEDER, UE), as well as the European Union’s Horizon 2020 research and innovation program under the Marie Sklodowska-Curie grant agreement 705464 (“SCOOBA”) to OSS. This is an AntECO (SCAR) contribution.

## Supplementary material

Figure S1. Rarefaction curves present the relationship between the sampling effort and the microbiome OTU richness in *Dendrilla antarctica* (Dant), *Sphaerotylus antarcticus* (Sant), *Mycale acerata* (Mace), *Hemigellius pilosus* (Ec) and surrounding seawater (SW).

Figure S2. Total abundance of the microbiome OTUs recovered from sponges and seawater samples as a function of its prevalence and classified by phyla. Log scale in the x-axis. Discontinuous line indicates 5% prevalence.

Figure S3. Venn diagrams showing the unique and shared microbiome OTUs among hosts (A) and including seawater samples (B) defined at distance of 0.03 (i.e., 97% similarity). *Dendrilla antarctica* (Dant), *Sphaerotylus antarcticus* (Sant), *Mycale acerata* (Mace), *Hemigellius pilosus* (Hpil) and seawater (SW).

Figure S4. Venn diagrams showing the unique and shared clone OTUs among hosts (A) and seawater samples (B) from Deception Island defined at distance of 0.03 (i.e., 97% similarity). *Dendrilla antarctica* (Dant), *Sphaerotylus antarcticus* (Sant), *Mycale acerata* (Mace), *Hemigellius pilosus* (Hpil) and seawater (SW).

Figure S5. Total abundance of the microbiome OTUs recovered from sponge samples as a function of its prevalence and classified by phyla. Log scale in the x-axis. Discontinuous line indicates 5% prevalence.

Figure S6. Total abundance of the microbiome OTUs recovered from seawater samples as a function of its prevalence and classified by phyla. Log scale in the x-axis. Discontinuous line indicates 5% prevalence.

Table S1. Core microbiome defined at 0.1% relative abundance and 90% prevalence and variable microbiome defined at 0.1% relative abundance and 10-90% prevalence. Abundances across samples are also shown at 0.1% and 1% (in some cases) relative abundances. OTUs in bold represent those with a minimum relative abundance of 1% that were found in 90% of the samples. Sources: *Dendrilla antarctica, Sphaerotylus antarcticus, Mycale acerata, Hemigellius pilosus* and seawater.

Table S2. Significantly different abundant OTUs in multiple comparisons among sources according to the false discovery rate (FDR) probabilities. Mean sequence count for the corresponding source is provided with colored values representing higher counts than the other sources compared. The taxonomy affiliation of each OTU is also shown with the percentage identity in parenthesis. Sources: sponges, seawater, *Dendrilla antarctica, Sphaerotylus antarcticus, Mycale acerata, Hemigellius pilosus*). Representing sponge Core Microbiome OTUs in bold (at 0.1% Relative Abundance and 90% Prevalence) and Variable OTUs in gray (at 0.1% Relative Abundance and 10-90% Prevalence).

File S1. Local blast results of the microbiome OTUs from *Dendrilla antarctica, Sphaerotylus antarcticus, Mycale acerata, Hemigellius pilosus* and seawater against the Sponge Earth Microbiome Project database. First hit, alignment matches and sequence identities are shown. Percentage of microbiome OTUs above identity thresholds is also shown. Sponge core OTUs are represented in bold and variable OTUs in gray.

File S2. Local blast results of the clone OTUs from *Dendrilla antarctica, Sphaerotylus antarcticus, Mycale acerata, Hemigellius pilosus* and seawater against the microbiome dataset from this study. First and second hit, alignment matches and sequence identities are shown. Percentage of clone OTUs above identity thresholds is also shown.

## References

Angulo-Preckler C, Cid C, Oliva F, Avila C (2015) Antifouling activity in some benthic Antarctic invertebrates by *in situ* experiments at Deception Island, Antarctica. Mar Environ Res 105:30–8.

Angulo-Preckler C, Figuerola B, Núñez-Pons L, Moles J, Martín-Martín R, Rull-Lluch J, Gómez-Garreta A, Avila C (2018) Macrobenthic patterns at the shallow marine waters in the caldera of the active volcano of Deception Island, Antarctica. Cont Shelf Res 157:20–31.

Apprill A, McNally S, Parsons R, Weber L (2015) Minor revision to V4 region SSU rRNA 806R gene primer greatly increases detection of SAR11 bacterioplankton. Aquat Microb Ecol 75:129–137.

Bavestrello G, Arillo A, Calcinai B, Cattaneo-Vietti R, Cerrano C, Gaino E, Penna A, Sara M (2000) Parasitic diatoms inside antarctic sponges. Biol Bull 198:29–33.

Bell JJ (2008) The functional roles of marine sponges. Estuar Coast Shelf Sci 79:341–353.

Caporaso JG, Kuczynski J, Stombaugh J, Bittinger K, Bushman FD, Costello EK, Fierer N, Peña AG, Goodrich JK, Gordon JI, et al (2010) QIIME allows analysis of high-throughput community sequencing data. Nat Methods 7:335–336.

Caporaso JG, Lauber CL, Walters WA, Berg-Lyons D, Lozupone CA, Turnbaugh PJ, Fierer N, Knight R (2011) Global patterns of 16S rRNA diversity at a depth of millions of sequences per sample. Proc Natl Acad Sci 108:4516–4522.

Cárdenas CA, Font A, Steinert G, Rondon R, González-Aravena M (2019) Temporal Stability of Bacterial Communities in Antarctic Sponges. Front Microbiol 10:1–14.

Cárdenas CA, González-Aravena M, Font A, Hestetun JT, Hajdu E, Trefault N, Malmberg M, Bongcam-Rudloff E (2018) High similarity in the microbiota of cold-water sponges of the Genus Mycale from two different geographical areas. PeerJ 6:e4935.

Cardoso J, van Bleijswijk J, Witte H, van Duyl F (2013) Diversity and abundance of ammonia-oxidizing Archaea and Bacteria in tropical and cold-water coral reef sponges. Aquat Microb Ecol 68:215–230.

Cerrano C, Arillo A, Bavestrello G, Calcinai B, Cattaneo-Vietti R, Penna A, Sarà M, Totti C (2000) Diatom invasion in the antarctic hexactinellid sponge Scolymastra joubini. Polar Biol 23:441–444.

Cerrano C, Calcinai B, Cucchiari E, Camillo C Di, Nigro M, Regoli F, Sarà A, Schiaparelli S, Totti C, Bavestrello G (2004) Are diatoms a food source for Antarctic sponges? Chem Ecol 20:57–64.

Cleary DFR, Swierts T, Coelho FJRC, Polónia ARM, Huang YM, Ferreira MRS, Putchakarn S, Carvalheiro L, van der Ent E, Ueng J, et al (2019) The sponge microbiome within the greater coral reef microbial metacommunity. Nat Commun 10:1644.

Drummond AJ, Ashton B, Buxton S, Cheung M, Cooper A, Duran C, Field M, Heled J KM & MS et al. (2010) G v5. 3. A at: http://www.geneious.co. (2010) Geneious v5.3. Available at: http://www.geneious.com.

Easson CG, Chaves-Fonnegra A, Thacker RW, Lopez J V. (2020) Host population genetics and biogeography structure the microbiome of the sponge Cliona delitrix. Ecol Evol ece3.6033.

Easson CG, Thacker RW (2014) Phylogenetic signal in the community structure of host-specific microbiomes of tropical marine sponges. Front Microbiol. doi: 10.3389/fmicb.2014.00532

Erwin PM, López-Legentil S, González-Pech R, Turon X (2012a) A specific mix of generalists: Bacterial symbionts in Mediterranean *Ircinia* spp. FEMS Microbiol Ecol 79:619–637.

Erwin PM, Pita L, López-Legentil S, Turon X (2012b) Stability of sponge-associated bacteria over large seasonal shifts in temperature and irradiance. Appl Environ Microbiol 78:7358–7368.

Glasl B, Smith CE, Bourne DG, Webster NS (2018) Exploring the diversity-stability paradigm using sponge microbial communities. Sci Rep 8:8425.

Henríquez M, Vergara K, Norambuena J, Beiza A, Maza F, Ubilla P, Araya I, Chávez R, San-Martín A, Darias J, et al (2014) Diversity of cultivable fungi associated with Antarctic marine sponges and screening for their antimicrobial, antitumoral and antioxidant potential. World J Microbiol Biotechnol 30:65–76.

Hentschel U, Hopke J, Horn M, Anja B, Wagner M, Hacker J, Bradley S, Friedrich AB, Moore BS (2002) Molecular evidence for a uniform microbial community in sponges from different oceans. Appl Environ Microbiol 68:4431–4440.

Hill MS, Sacristán-Soriano O (2017) Molecular and functional ecology of sponges and their microbial symbionts. In: Climate Change, Ocean Acidification and Sponges. Springer International Publishing, Cham, pp 105–142

Hooper JN a., Van Soest RWM (2004) Book Review: Systema Porifera. Guide to the classification of sponges. Invertebr Syst 18:233–234.

Jackson S a., Flemer B, McCann A, Kennedy J, Morrissey JP, O’Gara F, Dobson ADW (2013) Archaea appear to dominate the microbiome of Inflatella pellicula deep sea sponges. PLoS One 8:1–8.

Kennedy J, Flemer B, Jackson S a., Morrissey JP, O’Gara F, Dobson ADW (2014) Evidence of a putative deep sea specific microbiome in marine sponges. PLoS One. doi: 10.1371/journal.pone.0091092

Lee OO, Wang Y, Yang J, Lafi FF, Al-Suwailem A, Qian P-Y (2011) Pyrosequencing reveals highly diverse and species-specific microbial communities in sponges from the Red Sea. ISME J 5:650–664.

Lo Giudice A, Azzaro M, Schiaparelli S (2019) Microbial Symbionts of Antarctic Marine Benthic Invertebrates. In: The Ecological Role of Micro-organisms in the Antarctic Environment. pp 277–296

McClintock JB, Amsler CD, Baker BJ, van Soest RWM (2005) Ecology of Antarctic marine sponges: An overview. Integr Comp Biol 45:359–368.

Moitinho-Silva L, Bayer K, Cannistraci C V., Giles EC, Ryu T, Seridi L, Ravasi T, Hentschel U (2014) Specificity and transcriptional activity of microbiota associated with low and high microbial abundance sponges from the Red Sea. Mol Ecol 23:1348–1363.

Moitinho-Silva L, Nielsen S, Amir A, Gonzalez A, Ackermann GL, Cerrano C, Astudillo-Garcia C, Easson C, Sipkema D, Liu F, et al (2017a) The sponge microbiome project. Gigascience 6:1–7.

Moitinho-Silva L, Steinert G, Nielsen S, Hardoim CCP, Wu YC, McCormack GP, López-Legentil S, Marchant R, Webster N, Thomas T, et al (2017b) Predicting the HMA-LMA status in marine sponges by machine learning. Front Microbiol 8:1–14.

Moreno-Pino M, Cristi A, Gillooly JF, Trefault N (2020) Characterizing the microbiomes of Antarctic sponges: a functional metagenomic approach. Sci Rep 10:645.

Pita L, Erwin PM, Turon X, López-Legentil S (2013) Till death do us part: Stable sponge-bacteria associations under thermal and food shortage stresses. PLoS One. doi: 10.1371/journal.pone.0080307

Pita L, Rix L, Slaby BM, Franke A, Hentschel U (2018) The sponge holobiont in a changing ocean: from microbes to ecosystems. Microbiome 6:46.

Polónia ARM, Cleary DFR, Duarte LN, de Voogd NJ, Gomes NCM (2014) Composition of Archaea in Seawater, Sediment, and Sponges in the Kepulauan Seribu Reef System, Indonesia. Microb Ecol 67:553–567.

Polónia ARM, Cleary DFR, Freitas R, de Voogd NJ, Gomes NCM (2015) The putative functional ecology and distribution of archaeal communities in sponges, sediment and seawater in a coral reef environment. Mol Ecol 24:409–423.

Radax R, Hoffmann F, Rapp HT, Leininger S, Schleper C (2012) Ammonia-oxidizing archaea as main drivers of nitrification in cold-water sponges. Environ Microbiol 14:909–923.

Reveillaud J, Maignien L, Eren M a, Huber J a, Apprill A, Sogin ML, Vanreusel A (2014) Host-specificity among abundant and rare taxa in the sponge microbiome. ISME J 8:1198–209.

Rodríguez-Marconi S, De la Iglesia R, Díez B, Fonseca CA, Hajdu E, Trefault N (2015) Characterization of Bacterial, Archaeal and Eukaryote Symbionts from Antarctic Sponges Reveals a High Diversity at a Three-Domain Level and a Particular Signature for This Ecosystem. PLoS One 10:e0138837.

Sacristán-Soriano O, Winkler M, Erwin P, Weisz J, Harriott O, Heussler G, Bauer E, West Marsden B, Hill A, Hill M (2019) Ontogeny of symbiont community structure in two carotenoid-rich, viviparous marine sponges: comparison of microbiomes and analysis of culturable pigmented heterotrophic bacteria. Environ Microbiol Rep 11:249–261.

Schloss PD, Westcott SL, Ryabin T, Hall JR, Hartmann M, Hollister EB, Lesniewski RA, Oakley BB, Parks DH, Robinson CJ, et al (2009) Introducing mothur: Open-Source, Platform-Independent, Community-Supported Software for Describing and Comparing Microbial Communities. Appl Environ Microbiol 75:7537–7541.

Schmitt S, Deines P, Behnam F, Wagner M, Taylor MW (2011) Chloroflexi bacteria are more diverse, abundant, and similar in high than in low microbial abundance sponges. FEMS Microbiol Ecol 78:497–510.

Schmitt S, Tsai P, Bell J, Fromont J, Ilan M, Lindquist N, Perez T, Rodrigo A, Schupp PJ, Vacelet J, et al (2012) Assessing the complex sponge microbiota: core, variable and species-specific bacterial communities in marine sponges. ISME J 6:564–576.

Simister RL, Deines P, Botté ES, Webster NS, Taylor MW (2012) Sponge-specific clusters revisited: A comprehensive phylogeny of sponge-associated microorganisms. Environ Microbiol 14:517–524.

Steinert G, Wemheuer B, Janussen D, Erpenbeck D, Daniel R, Simon M, Brinkhoff T, Schupp PJ (2019) Prokaryotic Diversity and Community Patterns in Antarctic Continental Shelf Sponges. Front Mar Sci 6:1–15.

Taylor MW, Radax R, Steger D, Wagner M (2007) Sponge-associated microorganisms: evolution, ecology, and biotechnological potential. Microbiol Mol Biol Rev 71:295–347.

Thomas T, Moitinho-Silva L, Lurgi M, Björk JR, Easson C, Astudillo-García C, Olson JB, Erwin PM, López-Legentil S, Luter H, et al (2016) Diversity, structure and convergent evolution of the global sponge microbiome. Nat Commun 7:11870.

Turon M, Cáliz J, Garate L, Casamayor EO, Uriz MJ (2018) Showcasing the role of seawater in bacteria recruitment and microbiome stability in sponges. Sci Rep 8:15201.

Turon M, Uriz MJ (2020) New Insights Into the Archaeal Consortium of Tropical Sponges. Front Mar Sci 6:1–13.

Webster NS, Negri AP, Munro MMHG, Battershill CN (2004) Diverse microbial communities inhabit Antarctic sponges. Environ Microbiol 6:288–300.

Webster NS, Taylor MW (2012) Marine sponges and their microbial symbionts: Love and other relationships. Environ Microbiol 14:335–346.

Webster NS, Taylor MW, Behnam F, Lücker S, Rattei T, Whalan S, Horn M, Wagner M (2010) Deep sequencing reveals exceptional diversity and modes of transmission for bacterial sponge symbionts. Environ Microbiol 12:2070–2082.

Wilkins D, Lauro FM, Williams TJ, Demaere MZ, Brown M V., Hoffman JM, Andrews-Pfannkoch C, Mcquaid JB, Riddle MJ, Rintoul SR, et al (2013) Biogeographic partitioning of Southern Ocean microorganisms revealed by metagenomics. Environ Microbiol 15:1318–1333.

