## Supplementary figures and images for "Host species determine symbiotic community composition in Antarctic sponges (Porifera: Demospongiae)"

### Supplemental Figure S1

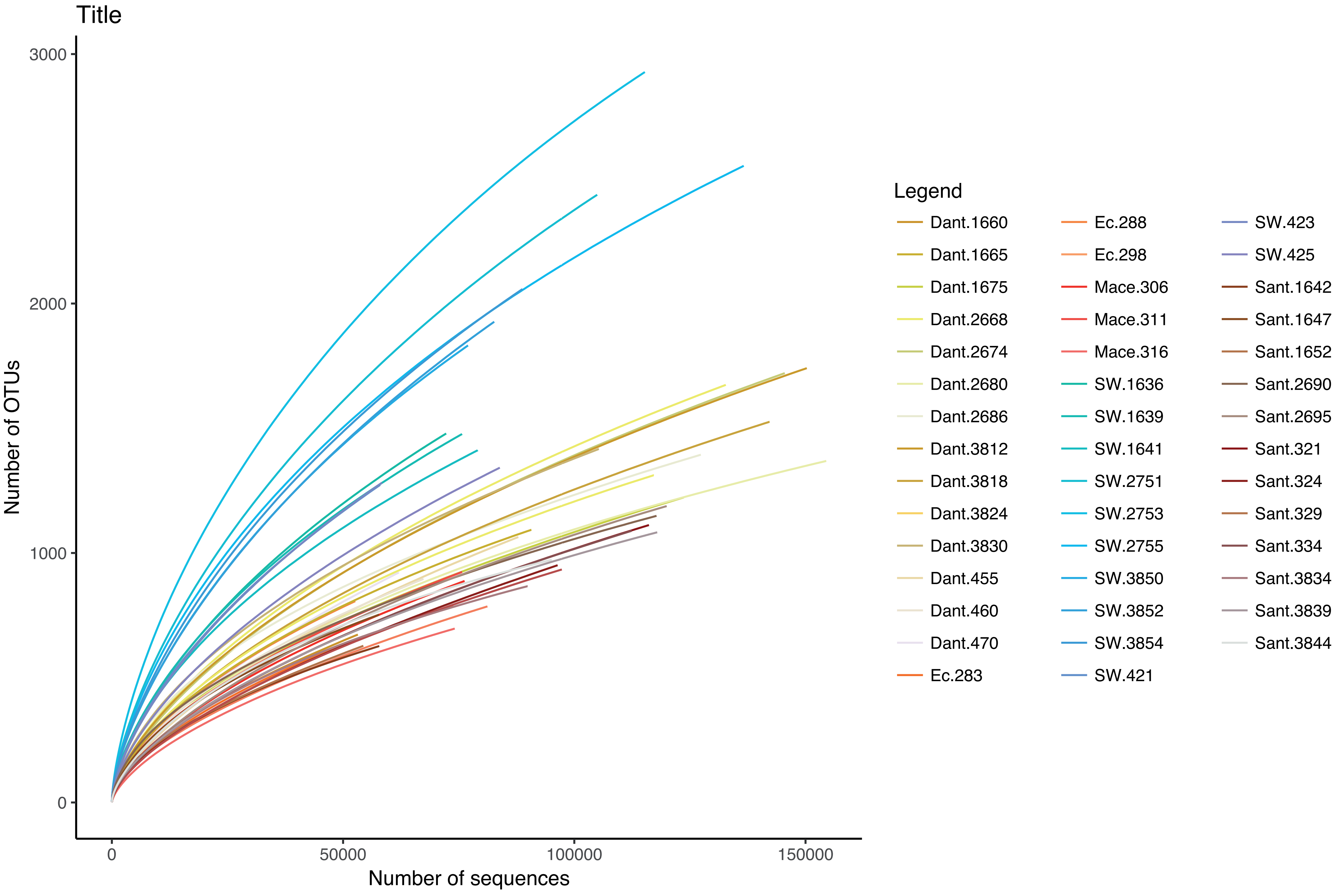

### Supplemental Figure S2

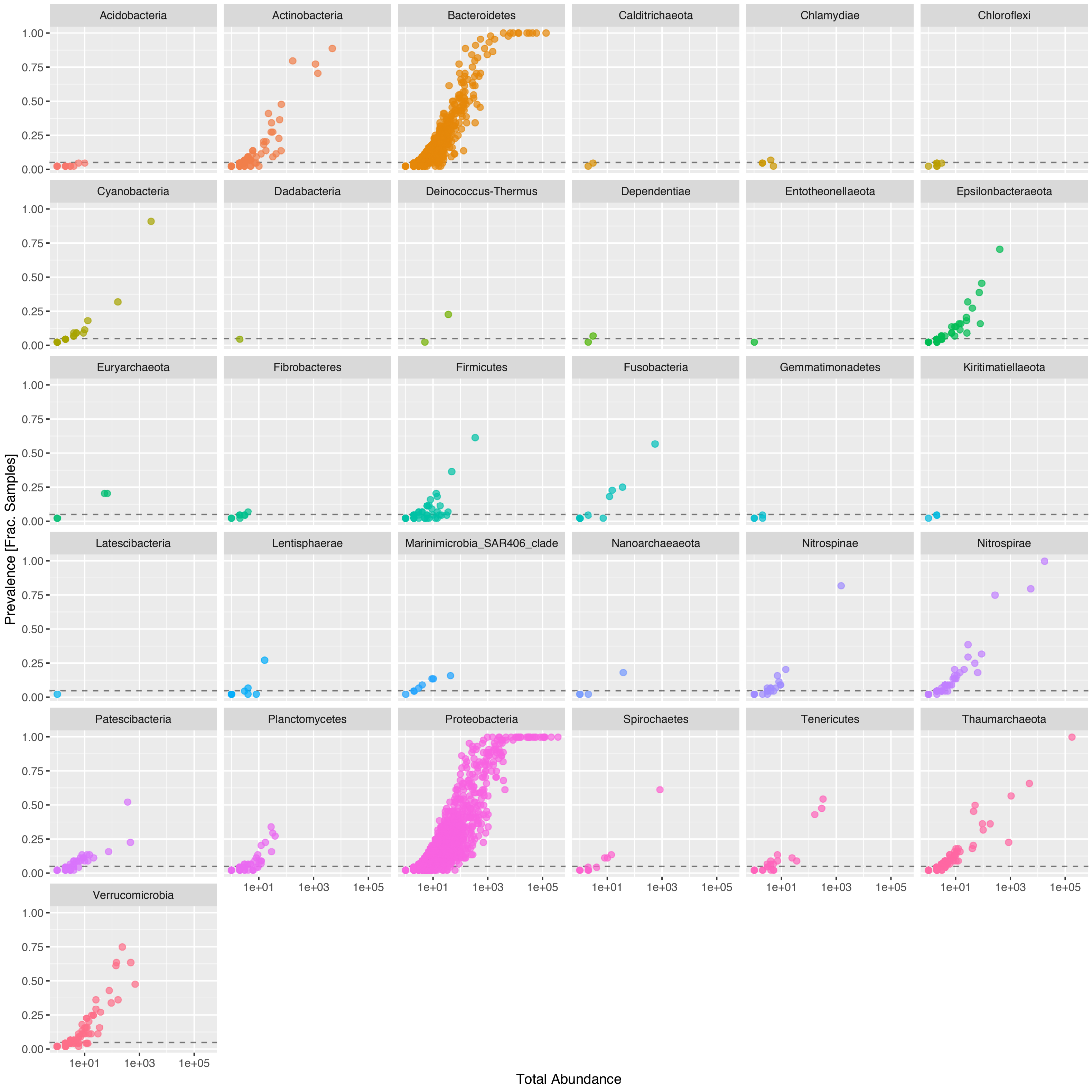

### Supplemental Figure S3

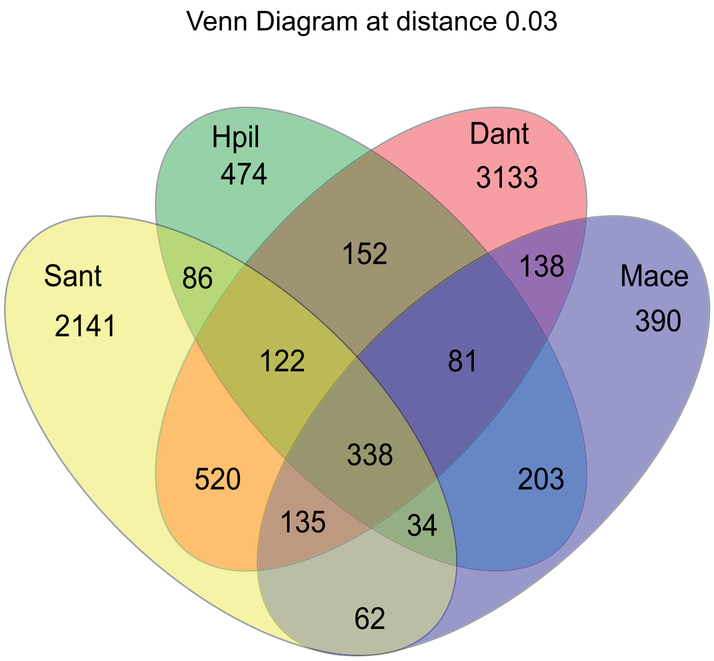

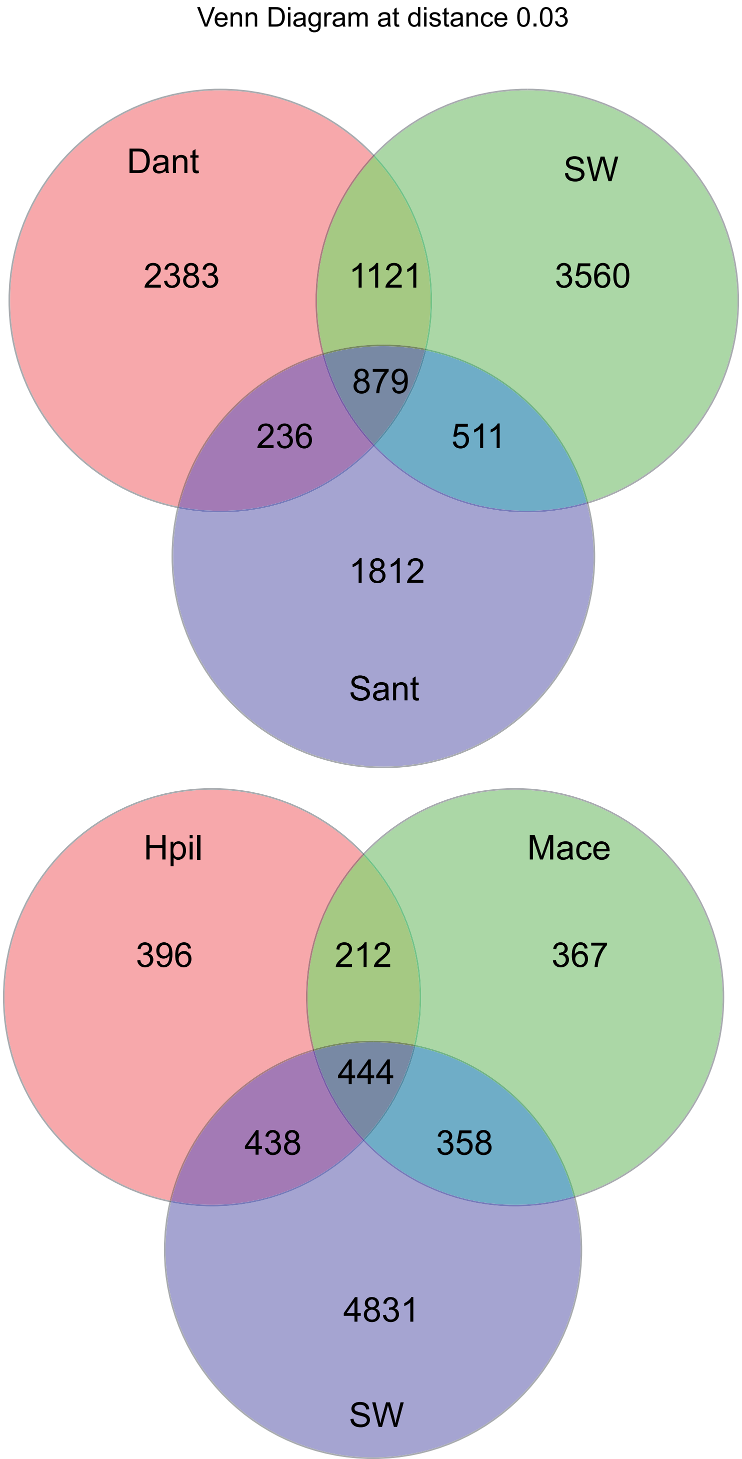

### Supplemental Figure S4

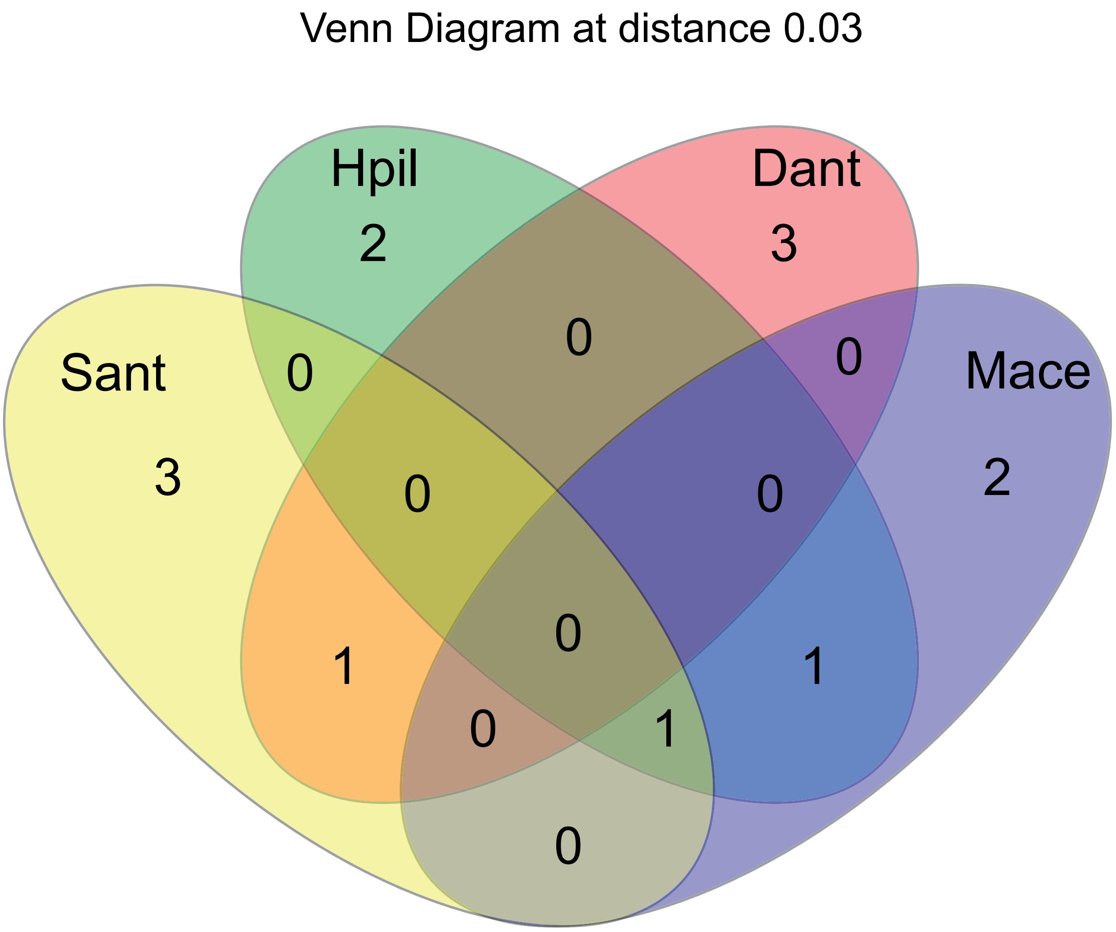

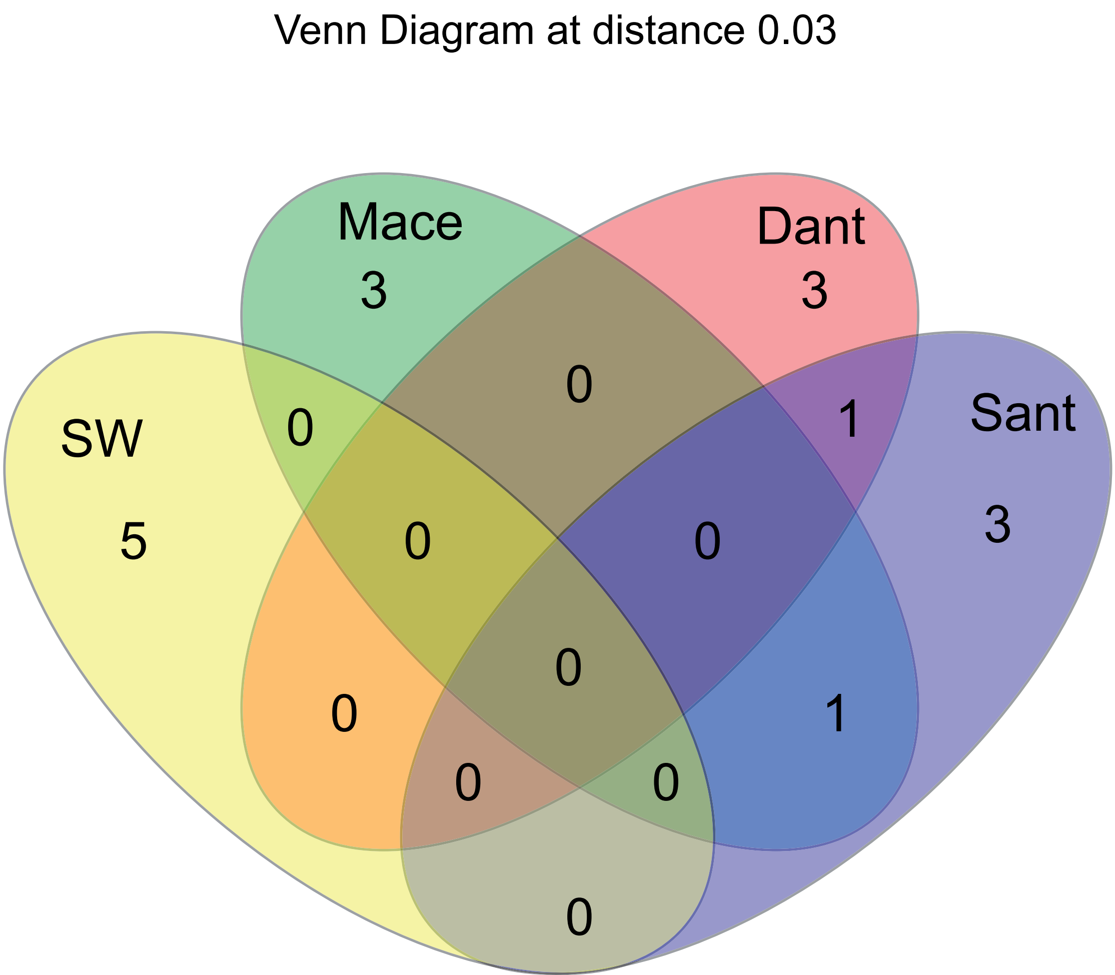

### Supplemental Figure S5

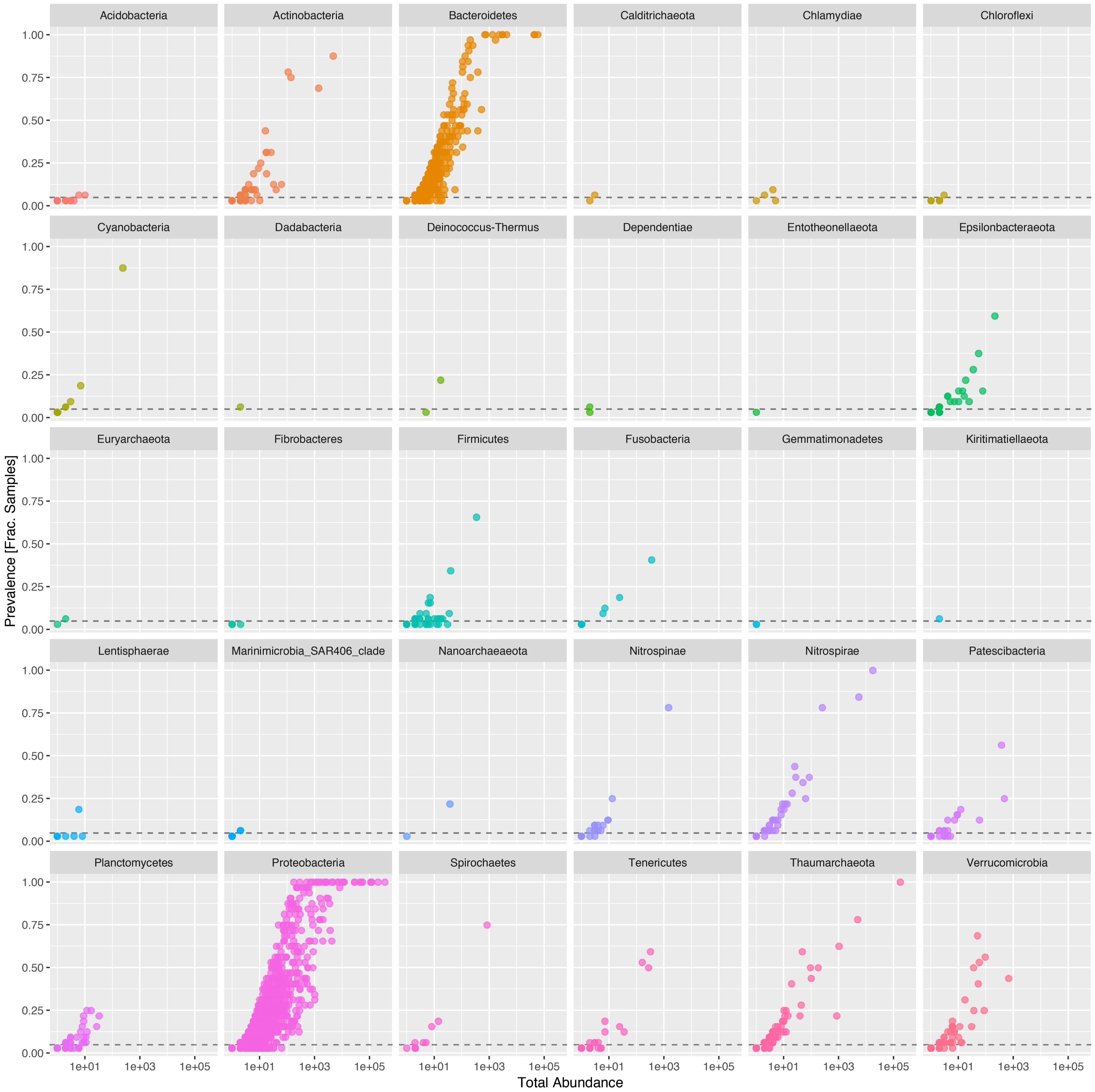

### Supplemental Figure S6

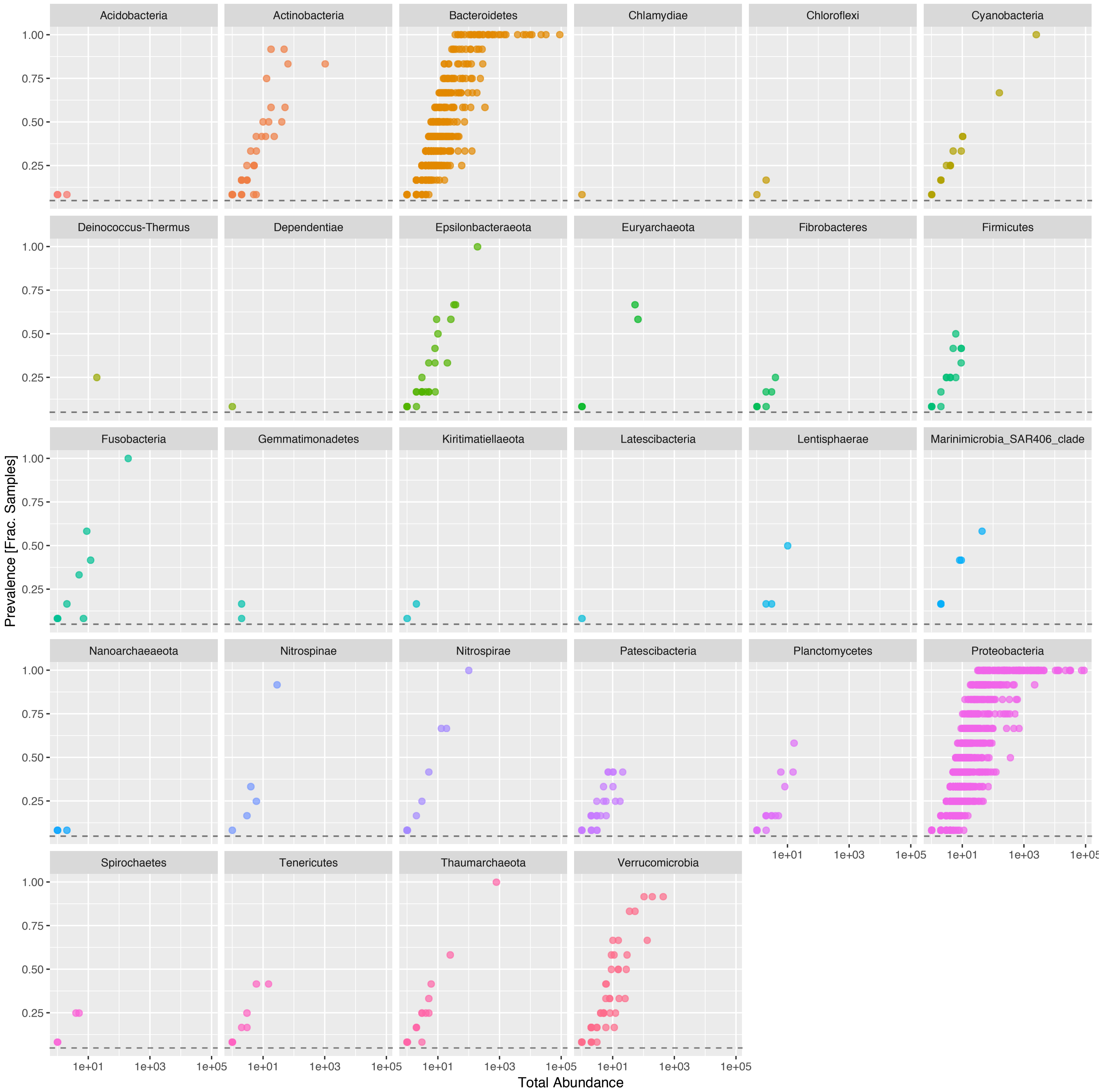
